# Direct evidence of abortive lytic infection-mediated establishment of Epstein-Barr virus latency during B-cell infection

**DOI:** 10.1101/2020.03.23.004440

**Authors:** Tomoki Inagaki, Yoshitaka Sato, Jumpei Ito, Mitsuaki Takaki, Yusuke Okuno, Masahiro Yaguchi, H. M. Abdullah Al Masud, Takahiro Watanabe, Kei Sato, Shingo Iwami, Takayuki Murata, Hiroshi Kimura

## Abstract

Viral infection induces dynamic changes in transcriptional profiles. Virus-induced and anti-viral responses are intertwined during the infection. Epstein-Barr virus (EBV) is a human gammaherpesvirus that provides a model of herpesvirus latency. To measure the transcriptome changes during the establishment of EBV latency, EBV-negative Akata cells were infected with EBV-EGFP and observed by transcriptome sequencing (RNA-seq) at 0, 2, 4, 7, 10, and 14 days after infection. We found transient downregulation of mitotic division-related genes, reflecting reprograming of cell growth by EBV. Moreover, a burst of viral lytic gene expression was detected in the early phase of infection. Experimental and mathematical investigations demonstrated that infectious virions were not produced in the pre-latent phase, suggesting the presence of an abortive lytic infection. Finally, we conducted fate mapping using recombinant EBV, enabling the noninvasive, continuous observation of infected cells during EBV infection. Our tracking analysis provided direct evidence that the abortive lytic infection in the pre-latent phase converges to latent infection during EBV infection of B-cells, shedding light on novel roles of viral lytic gene(s) in establishing latency.

**Author summary:** Viral infection is a complex process that activates both virus-triggered and host anti-viral responses. This process has classically been studied by snapshot analysis such as microarray and RNA-seq at discrete time points as population averages. Snapshot data lead to invaluable findings in host-pathogen interactions. However, these “snapshot” omics, even from a single cell, lack temporal resolution. Because the behavior of infected cells is highly dynamic and heterogenous, continuous analysis is required for deciphering the fate of infected cells during viral infection. Here, we exploited fate mapping techniques with recombinant Epstein-Barr virus (EBV) to track the infected cells and recorded a log of lytic gene expression during EBV infection. Our continuous observation of infected cells revealed that EBV established latency in B-cells via an abortive lytic infection in the pre-latent phase.

## Introduction

Numerous signaling events are triggered during the first few days of viral infection. Virus entry into the target cells results in the activation of cellular signaling pathways [1]. Concomitantly, viral pathogens are recognized by host sensor molecules, leading to activation of immune responses [2, 3]. Viruses rewire and modulate this interspecies interaction to meet their own needs, and consequently establish latency in their host cells.

Epstein-Barr virus (EBV), a gammaherpesvirus, is a widely dispersed enveloped virus that infects >90% of adults worldwide. It is associated with several types of human malignancies, with an incidence of 200,000 EBV-related cancers estimated annually [4]. Although most primary EBV infections are asymptomatic, EBV infection can cause infectious mononucleosis, especially when primary infection is delayed until late adolescence or early adulthood [5]. EBV establishes latent infection primarily in B-cells and typically persists for the life of the individuals [6, 7], although the virus can demonstrate both latent and lytic cycles in lymphocytes after primary infection. Thus, EBV provides a model system for studying how viruses, and particularly herpesviruses, establish latency in the cells.

In the latent state, the EBV genome is maintained as circular plasmids forming nucleosomal structures with histones, which expresses a limited number of viral gene products [8]. Therefore, no production of virus particles occurs during latent infection. Periodically, latent EBV switches from the latent stage into the lytic cycle to produce progeny viruses within its host cell. During the lytic infection, all EBV genes are expressed in a strictly regulated temporal cascade, and the circular genomes are amplified by the viral replication machinery, generating infectious virions [9, 10].

Accumulating evidence shows the abortive lytic cycle, as a third state of EBV infection occurring in the pre-latent phase of EBV primary infection [11–16] and within EBV-associated tumors [17–20]. In this state, the full lytic program is not induced due to an incomplete expression cascade of viral lytic genes; thus, infectious particles are not produced. Abortive lytic replication and its associated viral gene expressions are implicated in the pathogenesis of EBV-associated malignancies [17, 21, 22]. Transient lytic gene expressions in newly infected B-cells is essential for the emergence of lymphoblastoid cell lines [23].

mRNA expression profiling by RNA sequencing (RNA-seq) and qPCR analysis has provided important information on viral gene expression throughout these states of EBV-infected cells. However, such “snapshot” data lack temporal resolution, rendering them unsuitable to address the fate of infected cells during EBV infection. Continuous analysis is crucial to decoding this dynamic and heterogenous process. In this study, we address the fate of infected cells, which exhibit an abortive lytic infection, during EBV infection and recorded lytic gene expression by fate mapping with recombinant EBV. Our findings revealed that EBV is able to establish latency in B-cells via abortive lytic infection in the pre-latent phase.

## Results

### Transient burst of viral lytic gene expression during the pre-latent phase of EBV infection

To elucidate the gene expression dynamics during the course of EBV infection, we performed a time-course transcriptome analysis on EBV-infected cells (Figure 1A). Here, we used Akata cells, the Burkitt lymphoma cell line instead of primary B-cells, because we focus on the infection-mediated transcriptional changes during EBV infection without EBV-driven transformation and further subsequent analysis using genetics. EBV-negative Akata(-) cells were infected with EBV-EGFP, and infected cells were collected by FACS sorting at 2 days post-infection (dpi) according to EGFP-positivity. Subsequently, transcriptome information of the infected cells was obtained at 2, 4, 7, 10, and 14 dpi by mRNA-seq. Clustering analysis grouped genes into 10 clusters (Figure 1B), according to the temporal expression patterns across time points (Figure 1C). Gene ontology (GO) enrichment analysis was performed to interpret respective gene clusters (Figure 1C).

**Figure 1:**
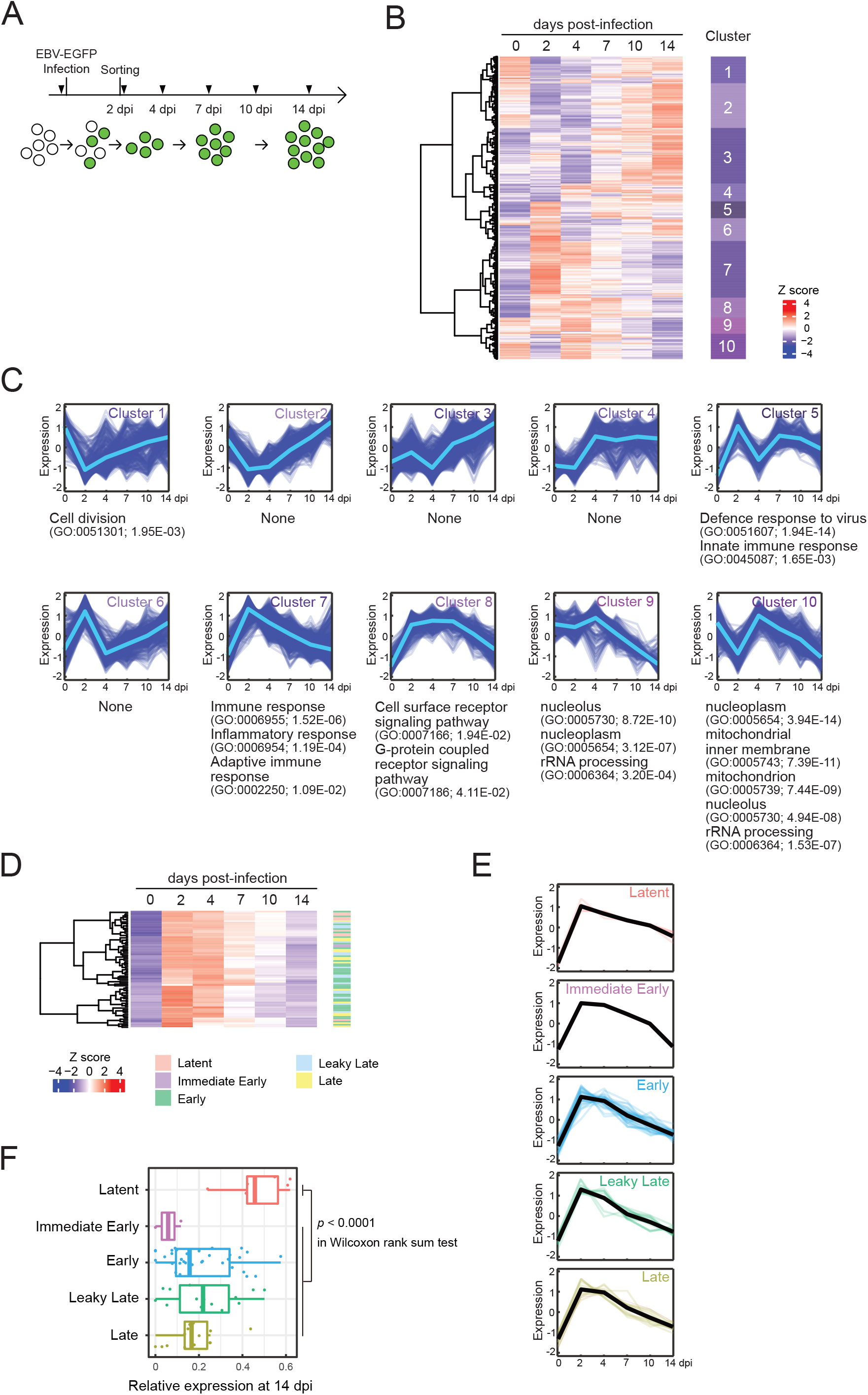
Transient burst of viral lytic gene expression in the pre-latent phase of EBV infection of B-cells. (A) Workflow of sample collection for RNA-seq analysis. Akata(-) cells were infected with EBV-EGFP. The infected cells were collected by FACS sorting at 2 dpi, and a portion of infected cells was harvested at the indicated time points (arrowheads). (B) Heatmap showing gene expression changes during EBV infection of Akata(-) cells. The 5,000 most diversely expressed genes of both the host and virus are included. Color indicates the normalized expression level (Z score). Gene clusters are indicated on the right side of the heatmap. (C) Temporal changes of gene expression in each cluster. The changes of respective genes (blue) and the mean value (light blue) are plotted. GO enrichment analysis was performed in each cluster, and the representative results (GO terms, GO Ids, and adjusted p-values) are shown on the right side of the graph. (D) Heatmap showing the viral gene expression changes during EBV infection of Akata(-) cells. (E) Temporal change of viral gene expression in each kinetic lytic gene. The changes of respective genes and the mean values (black) are plotted. Viral gene expression kinetics are categorized into five groups: latent, immediate early, early, leaky late, and late [47]. (F) Relative viral gene expression of latent, immediate early, early, leaky late, and late kinetics at 14 dpi.

Cluster 1 displayed an immediate decrease of expression followed by recovery of this decreased expression. Likewise, cluster 10 expression levels immediately decreased then rapid increased. These clusters preferentially included genes involving cell division and energy- or bio-syntheses (i.e., mitochondrion and rRNA processing-related genes), suggesting that cell growth was suppressed after viral infection. This observation is consistent with previous findings that rapid growth of EBV-infected cells is coupled with an increase in biomass for energy and production of biosynthetic intermediates [24]. Hammerschmidt and colleagues also showed that infected cells did not divide within the first three days of infection but rapidly re-commenced growth at 4 dpi, during EBV infection to naïve human B-cells [25]. Thus, EBV reprogrammed the transcriptome of infected cells during the initial stage of infection even without immortalization, similar to EBV infection of naïve or resting B-cells [15, 25, 26].

In agreement with data from EBV infection of human primary resting B-cells [15], clusters 5 and 7 displayed peak expression at 2 dpi enriched with genes involved in immune responses,

In parallel with cellular gene expression, viral genes are expressed in a dynamic but regulated manner during *de novo* infection of B-cells. Two days after infection, almost all viral genes including lytic genes were detected, and most of these genes had disappeared by 7 dpi (Figure 1D and E). Consequently, the pattern of viral gene expression demonstrated latent infection at 14 dpi (Figure 1F).

### Infectious virion production did not occur at the pre-latent phase

Upon EBV *de novo* infection, the synthesis of progeny virus was not observed in a previous study [16]. Our present study also confirmed this phenomenon (Figure 2A). Akata(-) cells were infected with EBV and washed after 2 h in PBS to remove unbound EBV inoculum. The cells were maintained and monitored for 7 dpi. Although infected cells were observed at 7 dpi, infectious virions were not detected in the supernatant at this time (Figure 2A).

**Figure 2:**
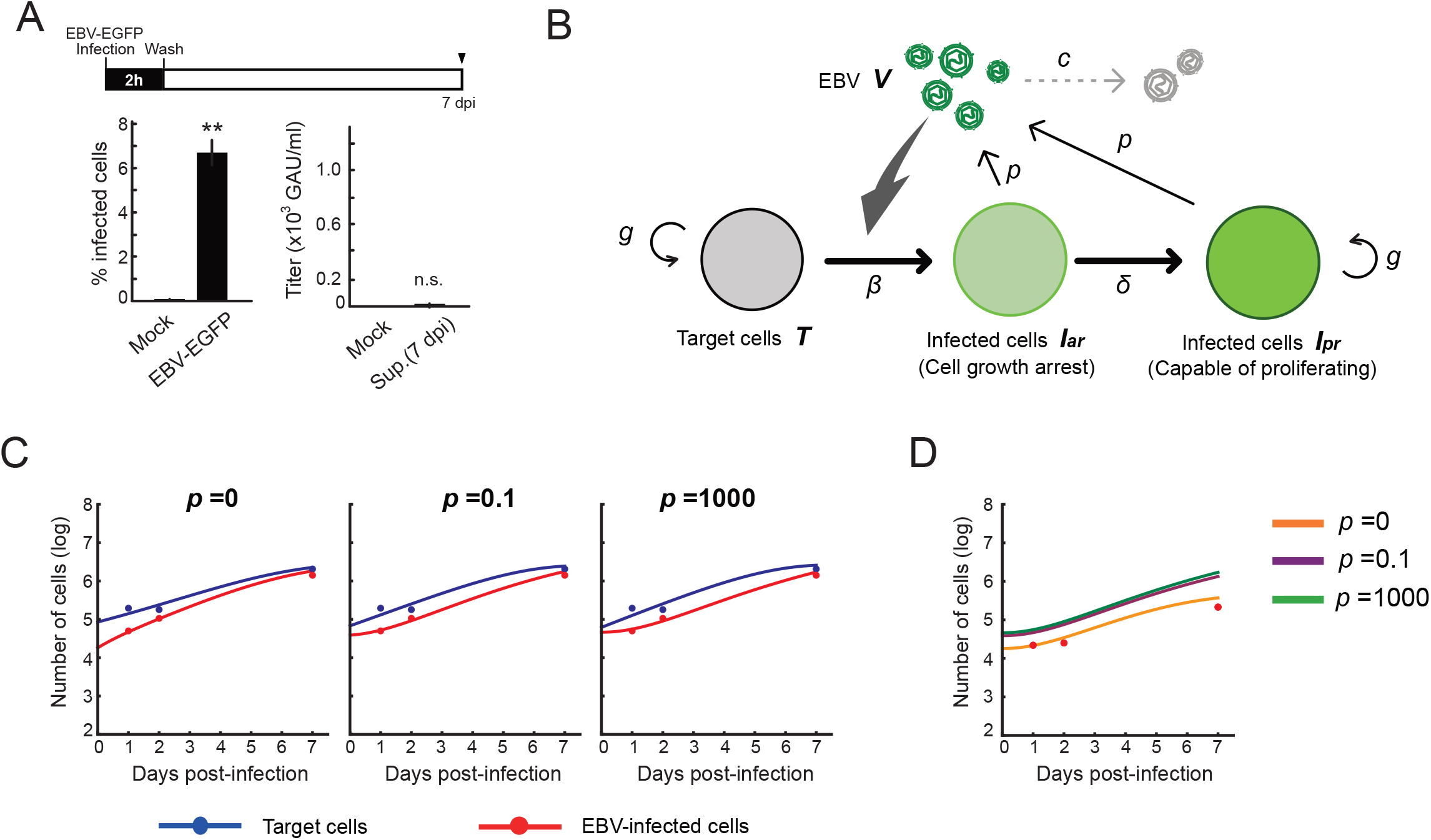
Experimental-mathematical investigation of progeny production during the pre-latent phase of EBV infection. (A) Infectious progeny virus is not detected from the culture supernatant during the pre-latent infection. Akata(-) cells were incubated with EBV-EGFP at room temperature for 2 h with agitation and then extensively washed with PBS to remove unbound virus. Cells and culture supernatant were harvested at 7 dpi. Infected cells were quantified by FACS (*left*). Virus titer in the supernatant was determined as described previously [46] (*right*). Results shown are the means ± SDs of three independent experiments. Double asterisks (**) indicate p < 0.01; n.s., not significant; dpi, days post-infection. (B) Mathematical model for EBV infection of B cells is described. The parameter *g* is the growth rate of cells, *K* is the carrying capacity, *β* is the cell-free infection rate, *δ* is the reciprocal number of cell growth arrest period, *p* is the progeny virus production rate, and *c* is the decay rate of EBV. Note that *p* = 0 corresponds with no progeny virus production. (C) With fixed values of *p* = 0,0.1, and 1000, we fitted Eqs. (3-6) to the experimental data and describe the solid curves of the best-fit solution. (D) The number of infected cells were calculated from mathematical model with estimated parameters. Except in the case of *p* = 0, prediction by the mathematical model was unable to reproduce the EBV infection dynamics properly.

We next carried out an *in silico* simulation based on a mathematical model and estimated parameters from experiments. Several studies on other viruses have established mathematical models that describe cell-free infection [27, 28]. We constructed a model for EBV infection considering two opposite assumptions of the existence or absence of progeny production, and we estimated parameters by fitting our experimental data without washing out unbound virus (see **mathematical modeling** in ***Materials and Methods***, and Figure 2B and C). Using these estimated parameters, we calculated EBV infection dynamics *in silico* under the assumption of washing out unbound virus, then compared our mathematical prediction and our experimental data under parallel conditions. Interestingly, as shown in Figure 2D, our model accurately described the actual EBV infection dynamics best in the case of *p* = 0, suggesting no progeny production in the pre-latent phase. Taken together, the transient expression of lytic genes during the pre-latent phase reflected an abortive lytic infection of EBV.

### Abortive lytic infection in the pre-latent phase transitioned to latent infection

RNA-seq analysis at discrete time points during EBV infection showed the abortive lytic infection in the pre-latent phase, consistent with other studies [11, 16]. However, because snapshot analyses such as RNA-seq lack temporal resolution, it remains unclear whether the cells showing abortive lytic infection during pre-latent phase are able to shift into latently infected cells.

We thus applied fate mapping techniques using the Flippase (Flpe) recombinase-flippase recognition target (FRT) system [29] to trace the fate of cells exhibiting abortive lytic infection in the pre-latent phase of EBV infection. Reporter cells were generated and isolated form Akata(-) cells, which transduced a red fluorescent protein (*DsRed*) reporter flanked by a neomycin-resistant gene containing STOP codon (*FRT-STOP)-FRT* sequence (Figure 3A). In cells expressing both Flpe and the reporter, Flpe specifically activated the reporter by excising the STOP sequence. Indeed, we confirmed the DsRed expression in the reporter cells in the presence of Flpe (Figure 3B). Notably, however, a few reporter cells expressed DsRed without Flpe expression. Furthermore, we generated the recombinant EBV(BMRF1p-Flpe), which expresses Flpe under the control of the BMRF1 promoter (Figure 3C). The *BMRF1* gene is categorized as an early lytic gene and encodes the DNA polymerase processivity factor, which associates with the polymerase catalytic subunit to enhance the polymerase processivity and exonuclease activity [30]. The BMRF1 protein is also known as early antigen diffused (EA-D) and is used as a clinical marker for EBV infection [31, 32]. Because the BMRF1 protein is abundantly expressed during lytic replication [33], we chose the BMRF1 promoter for monitoring the EBV lytic gene expression. Infectious virus particles of EBV(BMRF1p-Flpe) were prepared by transient trans-complementation of BMRF1.

**Figure 3:**
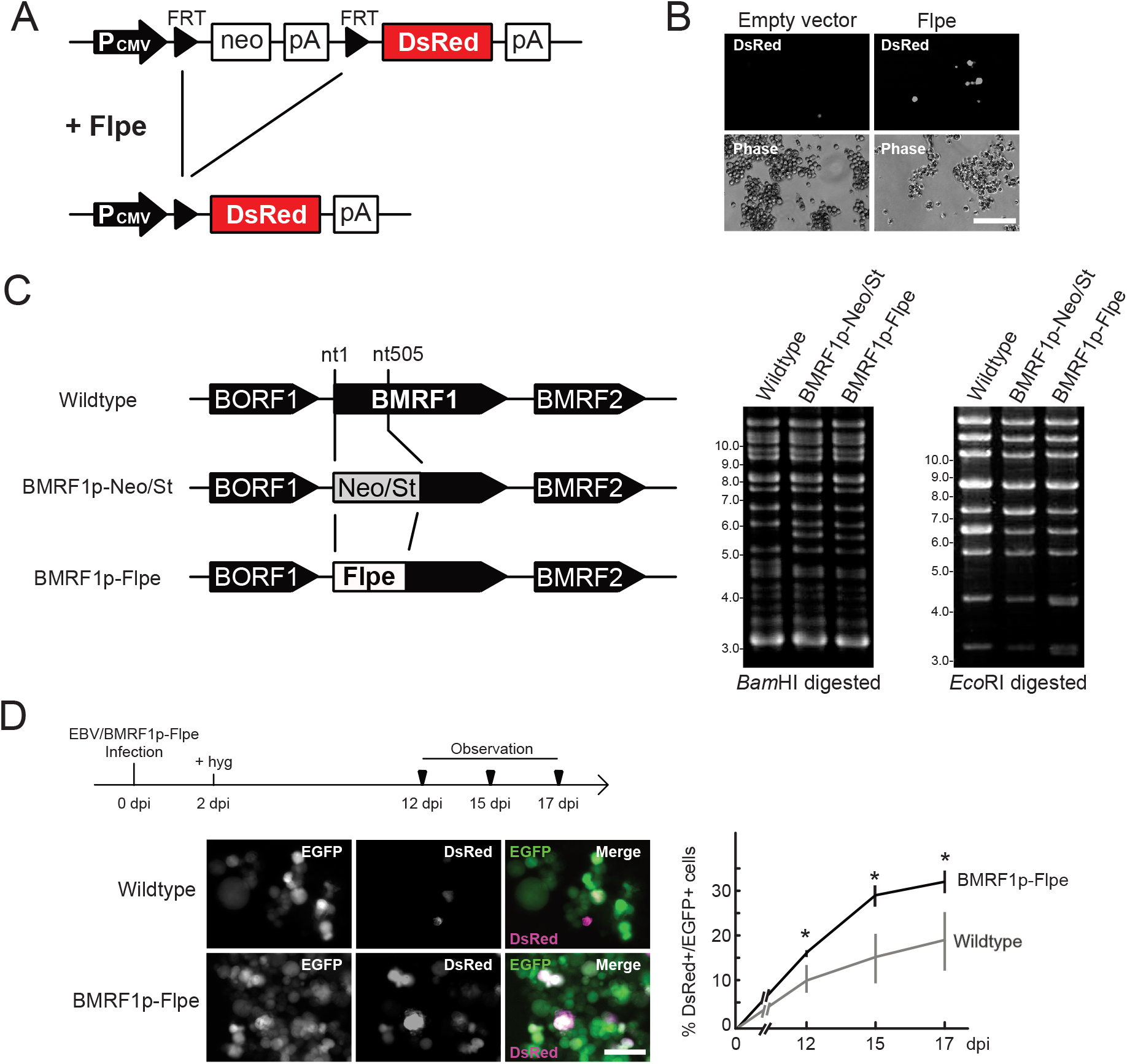
EBV establishes latency in B-cells via an abortive lytic infection in the pre-latent phase. (A) Schematic representation of the genetic elements in the Flpe-FRT system. Flpe recombinase can recombine FRT sites in the ubiquitously expressed reporter construct to remove the STOP (neomycin resistant gene and polyA signal; neo-pA) cassette. Upon removal of this STOP, the reporter DsRed is expressed in the cells and all their progeny. (B) DsRed was expressed in the presence of Flpe. Akata/FNF-DsRed reporter cells were transfected with a Flpe expression plasmid. Scale bar; 100 μm. (C) Schematic diagram of recombinant EBV (EBV/BMRF1p-Flpe) construction. The Neo/St cassette, containing neomycin resistance and streptomycin sensitivity genes, was inserted between nucleotide 1 and 505 of the BMRF1 gene to prepare an intermediate, and this was replaced with the Flpe sequence to construct the EBV/BMRF1p-Flpe, which expresses Flpe under the control of the BMRF1 promoter (*left*). Successful recombination was confirmed by the electrophoresis of the EBV-BAC after *Bam*HI and *Eco*RI digestion (*right*). (D) Continuous tracing of abortive lytic cells with recombinant EBV. Akata/FNF-DsRed cells were infected with EBV/BMRF1p-Flpe. Two days after infection, hygromycin was added to the medium to select infected cells. Infected cells were continuously observed (arrow heads) until 17 dpi. The number of DsRed-positive and EGFP-positive cells was counted at indicated time points. Results shown are the means ± SDs of three independent experiments. Images were obtained at 17 dpi. Asterisk (*) indicates p < 0.05; dpi, days post-infection. Scale bar; 50 μm.

Using Akata/FNF-DsRed reporter cells and recombinant EBV, fate mapping of infected cells was performed. The reporter cells were infected with EBV(BMRF1p-Flpe) and were continuously observed during the pre-latent phase. At 17 dpi, approximately 30% of infected cells, where EBV have established latency, expressed DsRed (Figure 3D). This suggests a history of lytic gene expression in these cells during the pre-latent phase of EBV infection. Therefore, we found that the abortive lytic infection of EBV transitioned to latent infection during *de novo* infection of B-cells with EBV.

## Discussion

EBV infects resting B-cells and transforms them into lymphoblastoid cell lines (LCLs) *in vitro*. LCLs share many common features with posttransplant lymphoproliferative disease and AIDS lymphomas [34]. Thus, LCLs have been extensively used to study the mechanisms by which EBV causes cancers [15, 35–37]. However, because dynamic changes in viral infection overlap with the process involved in EBV-caused transformation during infection of resting B-cells, transcriptomic changes between these two processes are indistinguishable. Here, we used EBV-negative cell lines derived from EBV-positive Burkitt lymphoma to evaluate the effect of EBV infection on host transcription in a time-course study. Using the Akata(-) cell line in this study enabled us to manipulate its genome for transducing our reporter system, as discuss below. Similar to previous studies using primary B-cells [15, 26], our RNA-seq analysis on Akata(-) cells with EBV infection showed transient cell growth arrest for 2 dpi and re-proliferation thereafter (Figure 1C). These findings suggest that post-infectious growth arrest is associated with EBV infection, but not with EBV-mediated transformation. Of note is that this transient growth arrest after infection was additionally reflected in our mathematical model of EBV infection (see **mathematical modeling** in ***Materials and Methods***).

Viral lytic genes were expressed transiently in newly infected cells during the pre-latent phase of EBV infection (Figure 1D). Because EBV does not initiate the *de novo* synthesis of virus progeny upon primary infection, expression of these viral genes indicate an abortive lytic infection. However, comprehensive analyses of progeny production during the pre-latent phase have not been elucidated to date. In this study, our *in silico* simulation of EBV infection supported a lack of progeny virus production. Under the assumption that even a small amount of progeny virus is produced, the number of infected cells derived from our model were unable to trace the experimental data (Figure 2B-D), providing a theoretical evidence in support of our experimental observations as well as previous studies [16, 38].

RNA-seq data is a snapshot at a single time point and lacks temporal resolution. Thus, these data miss the exact timing and order of events underlying infection. Continuous observation with cell tracking overcomes this obstacle. Although low sensitivity of the reporter is one of the limitations in this study (Figure 3B), our fate mapping system with recombinant EBV successfully revealed that some cells display a record of lytic gene expression among the cells where EBV established latency (Figure 3D). This finding suggests that the transient expression of lytic genes contributes to the establishment of latent infection in infected cells. In accordance with this scenario, it has been reported that viral proteins BNLF2a and vIL-10/BCRF1 are expressed transiently upon the pre-latent phase of infection, and they prevent immune recognition and elimination of newly EBV-infected cells [11].

In summary, combining the techniques of population-based averaged snapshot analysis and a continuous tracking system with fluorescent protein expression, we demonstrated herein that EBV established latency in B-cells via an abortive lytic infection in the pre-latent phase, implying the reversibility of the abortive lytic state during EBV infection. Our findings shed light on novel roles of EBV lytic genes in the initial, pre-latent phase of B-cell infection.

## Materials and Methods

### Cells

Akata(-) cells [39] were cultured in RPMI1640 medium containing 10% fetal bovine serum (FBS). AGS/EBV-EGFP cells, which kindly provided by Dr. Hironori Yoshiyama [40], were grown in F-12 HAM’s medium supplemented with 10% FBS and 400 μg/mL G418. HEK293 (a kind gift from Dr. Henri-Jacques Delecluse), HEK293T (ATCC CRL-3216), and HEK293/EBV-Bac cells were maintained in DMEM supplemented with 10% FBS. All cells were maintained at 37°C in an atmosphere of 5% CO_2_.

### Plasmids and lentiviral vector

To construct the lentiviral reporter plasmid (CSII-CMV-FNF-DsRed), the FNF-DsRed fragment was generated by PCR from pCAFNF-DsRed, which was a kind gift from Dr. Connie Cepko (Addgene plasmid #13771), and was inserted into the *Bam*HI-*Xba*I site of the CSII-CMV-MCS-IRES2-Venus plasmid (generously gifted by Dr. Hiroyuki Miyoshi). The inserted DNA sequence was confirmed by direct DNA sequencing.

The lentiviral vector was prepared by recovering the culture supernatant of 293T cells transfected with CSII-CMV-FNF-DsRed together with expression plasmids for HIV-1 Gag-Pol and Rev (pCMVR8.74, Addgene plasmid #22036; kindly provided by Dr. Didier Trono), and for VSV-G (generously provided by Dr. Yasuo Ariumi).

The Flpe expression plasmid (pCSFLPe) was the kind gift of Dr. Gerhart Ryffel (Addgene plasmid #31130). pcDNA-BZLF1, pcDNA-gB and pcDNA-BMRF1 were previously described [41–43].

### Recombinant EBV-Bac

The original of EBV-BAC DNA (B95-8 strain) was kindly provided by Dr. Wolfgang Hammerschmidt [44]. To construct the BMRF1 promoter-driven Flpe expression EBV-Bac (EBV(BMRF1p-Flpe)), homologous recombination was undertaken in *E. coli* as described previously [45] with the oligonucleotide primers listed in Table 1. After construction of recombinant EBV-BAC strains, DNA was digested with *Bam*HI or *Eco*RI and resolved by agarose gel electrophoresis.

**Table 1:**
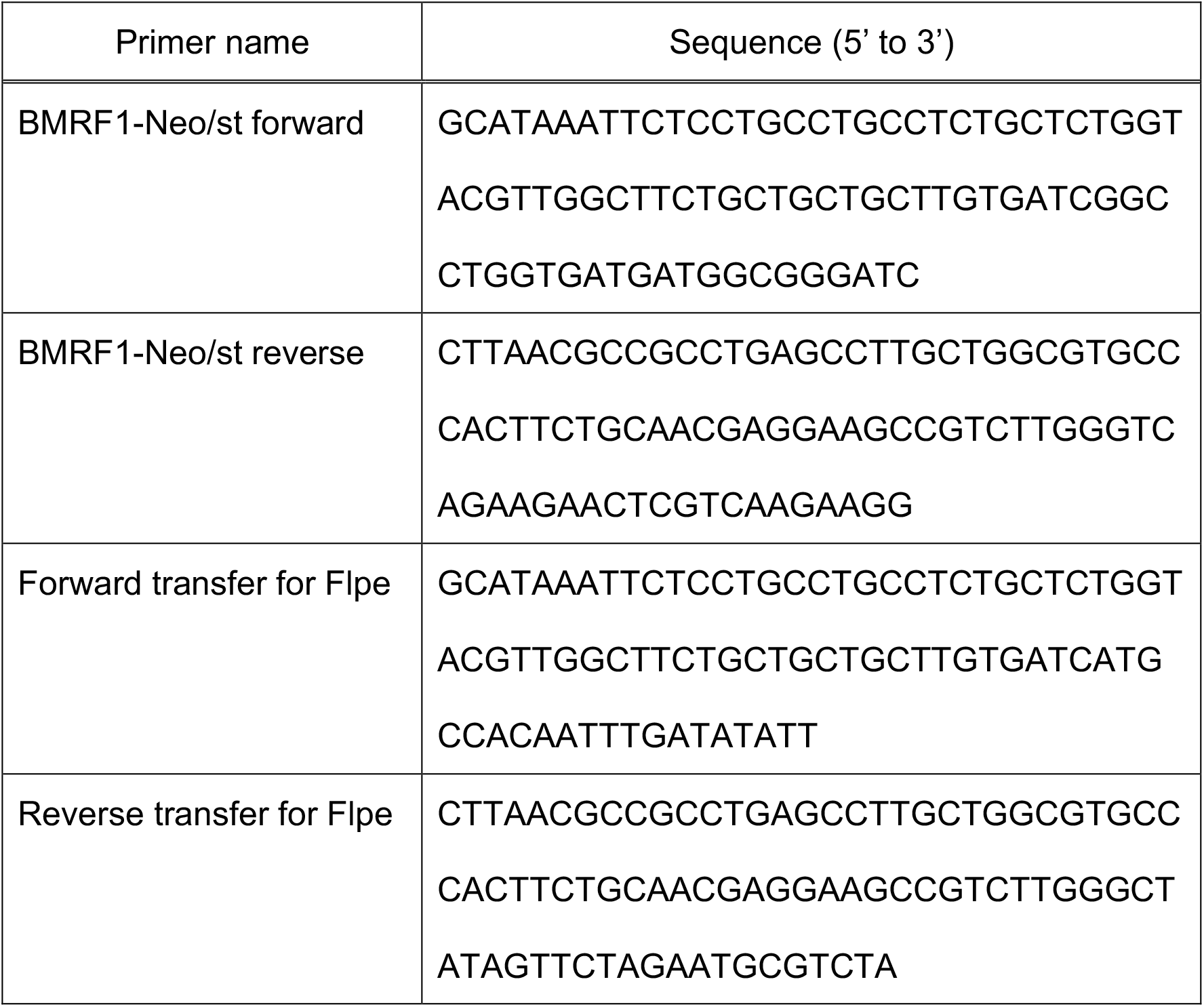
Oligonucleotide primers used for generation of recombinant EBV.

HEK293 cells were transfected with EBV-BAC DNA using Lipofectamine 2000 reagent (Thermo Fisher Scientific, Waltham, MA, USA) followed by hygromycin selection, and EGFP-positive cell colonies were selected for preparation of cell clones.

### EBV preparation

EGFP-EBV was obtained from the eight-day-old cell-free supernatant of AGS/EGFP-EBV cells. The cell-free supernatant was passed through 0.45 μm filters and then used as a virus stock.

For preparation of recombinant EBV, HEK293/EBV(BMRF1p-Flpe) and HEK293/EBV(WT) [46] cells were transfected with BZLF1 expression plasmid together with the gB expression plasmid using the Neon Transfection System (Thermo Fisher Scientific) to induce lytic replication. Cells and culture supernatants were collected, freeze-thawed, and centrifuged. Supernatant from the centrifugation was filtered and used as a virus stock.

### RNA-seq

Akata(-) cells were first spun down and resuspended in medium containing virus supernatant. Cells were incubated at room temperature for 2 h with agitation. The cells were spun again and resuspend in fresh medium. EBV-infected Akata cells that express EGFP were sorted by a FACS Aria II Cell Sorter (BD Biosciences, San Jose, CA, USA) at 2 dpi and then cultured. The cells were harvested for RNA preparation at later time points.

Total RNA was extracted using a Nucleospin RNA XS kit (Takara Bio, Kusatsu, Japan). Evaluation of RNA, RNA-seq library preparation, illumina sequencing, and data preprocessing were performed as described previously [17].

Gene expression levels were normalized as counts per million (CPM) followed by log2-transformation with a pseudo-count of 1. The 5,000 most diversely expressed viral and host genes were used for downstream analyses. After standardization, Euclid distances among genes were calculated according to their expression patterns. Clustering analysis of the genes was performed using the Ward method based on the Euclid distances. All analyses were performed on R (version 3.5.2).

GO enrichment analysis was performed according to an overlap-based method with one-sided Fisher’s exact test. The family-wise error rate (i.e., adjusted p-value) was calculated using the Holm method. As a source of gene sets, we used “GO biological process” and “GO cellular component” obtained from the Gene Ontology consortium (http://geneontology.org/; GO validation date: 08/30/2017).

### Data collection for mathematical modeling

Growth curves were measured as described previously [14]. Briefly, Akata(-) cells were seeded at a density of 1 × 10^5^ cells/mL. Every 48 h, the number of viable cells was counted. For EBV infection, 2 × 10^5^ cells of Akata(-) cells were first spun down and resuspended in 3 mL of medium containing EBV-EGFP (1 × 10^5^ green Akata units (GAU)). For washout conditions, 2 × 10^5^ cells of Akata(-) cells were incubated with EBV-EGFP (1 × 10^5^ GAU) at room temperature for 2 h with agitation. The infected cells were washed three times with 40 mL of PBS, and then suspended with 3mL of medium. The infected cells were seeded into a low-binding 35-mm dish (PrimeSurface dish MS-9035X; Sumitomo Bakelite, Tokyo, Japan) and maintained in a 37°C incubator at 5% CO_2_. The amount of virus particles in the culture supernatant and the number of infected cells were routinely measured as follows: a portion (400 μL) of the infected cell culture was routinely harvested, and the amount of released infectious virions in the culture supernatant was quantified as described previously [46]. The cell number was counted as described previously [14]. The percentage of infected cells was quantified by FACS [46].

For virus decay analysis, EBV-EGFP (1 × 10^5^ GAU) was suspended in 3mL of medium and incubated (37°C/5% CO_2_). Samples (100 μL) were routinely harvested, and the virus titer was quantified as described previously [46].

### Mathematical modeling

We independently estimated the growth kinetics of Akata cells by the following mathematical model (Eq. 1) from the separate experiments (see growth curve section in **Data collection for mathematical modeling**):

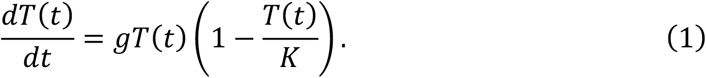

where the variable *T*(*t*) is the number of Akata cells, and the parameters *g* and *K* are the growth rate and the carrying capacity of the cell culture, respectively. We also estimated the viral decay kinetics by the following equation (Eq. 2) from the separate experiments (see virus decay section in **Data collection for mathematical modeling**):

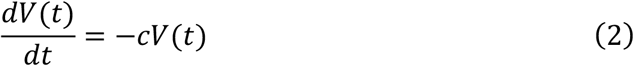

where *V*(*t*) is the number of virions, and *c* represents the viral decay rate. These parameters, *g*, *K* and *c*, are fixed hereafter.

Coupling EBV infection dynamics with Eqs.(1 and 2), we proposed the following mathematical models (Eqs. 3-6) considering two opposite assumptions of the presence or absence of progeny virus production:

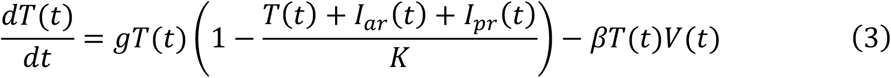

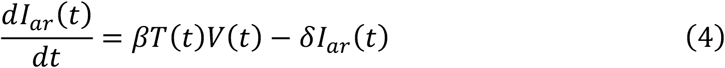

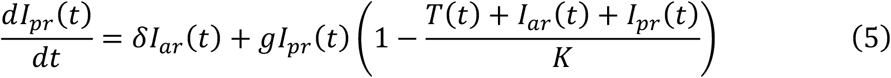

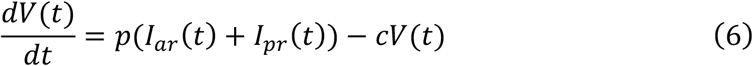

where *I_ar_*(*t*) is the number of EBV-infected cells with cell growth arrest, *I_pr_*(*t*) is that of all other EBV-infected cells, *β* is the infection rate, *δ* is the reciprocal number of cell growth arrest period (i.e., *δ* = 0.5 here), and *p* is the progeny virus production rate. Note that *p* = 0 corresponds with no virus progeny production. Initial values of the number of target cells and infected cells are given by *T*_0_ and *I*_*ar*,0_, respectively. In our data fitting, we estimated *β*, *T*_0_, and *I*_*ar*,0_ for *p* = 0,0.1,1000, fixing the estimated values of *g*, *K*, and *c* as described above. We summarized all estimated parameters in Table 2.

**Table 2:** Estimated parameters by fitting the mathematical model to experimental data.

| Symbol | Unit | Value |  |  |
| --- | --- | --- | --- | --- |
| | | $p = 0$ | $p = 0.1$ | $p = 1000$ |
| $g$ | day<sup>-1</sup> | $7.33 \times 10^{-1}$ | | |
| $K$ | cell | $6.51 \times 10^6$ | | |
| $c$ | day<sup>-1</sup> | $2.44 \times 10^{-1}$ | | |
| $\beta$ | (day×cell)<sup>-1</sup> | $2.46 \times 10^{-6}$ | $8.53 \times 10^4$ | $1.80 \times 10^4$ |
| $T_0$ | cell | $6.36 \times 10^{-7}$ | $6.76 \times 10^4$ | $3.88 \times 10^4$ |
| $I_{ar,0}$ | cell | $8.08 \times 10^{-11}$ | $6.17 \times 10^4$ | $4.62 \times 10^4$ |

### Fate mapping of EBV-infected cells

For generating reporter cells, Akata(-) cells were inoculated with the lentiviral vector harboring the CMV promoter-driven FNF-DsRed cassette and followed by G418 selection (750 μg/mL).

Akata/FNF-DsRed cells were incubated in medium containing EBV(BMRF1p-Flpe) at room temperature for 2h with agitation. The cells were spun down, resuspend in fresh medium, and then cultured. Expressions of fluorescent proteins, EGFP and DsRed were observed at discrete time points. Images were acquired using Zeiss Axio Observer microscope (Carl Zeiss, Oberkochen, Germany).

### Statistical analysis

Results are shown as the means ± standard deviation (SD) of three independent experiments. Statistical analyses were performed using Microsoft Excel and R (version 3.5.2). Unpaired Student’s *t*-test was used to determine significance between two groups. p-values <0.05 were considered significant.

### Data Availability

The RNA-seq raw data have been deposited in the DNA Data Bank of Japan (DDBJ; https://www.ddbj.nig.ac.jp/index-e.html) Sequence Reads Archive (DRA) under the accession number DRA009706.

## Acknowledgements

The authors thank Wolfgang Hammerschmidt, Henri-Jacques Delecluse, Hironori Yoshiyama, Connie Cepko, Gerhart Ryffel, Hiroyuki Miyoshi, Yasuo Ariumi and Didier Trono for providing invaluable materials; Shuko Kumagai, Mai Suganami, and Tomoko Kunogi for technical supports; and the Division for Medical Research Engineering at Nagoya University Graduate School of Medicine for technical support of cell sorting and next-generation sequencing.

